# On the importance of being stable ‒ evolutionarily frozen species can win in fluctuating environment

**DOI:** 10.1101/139030

**Authors:** Jaroslav Flegr, Petr Ponížil

## Abstract

The ability of organisms to adaptively respond to environmental changes (evolvability) is usually considered to be an important advantage in interspecific competition. It has been suggested, however, that evolvability could be a double-edged sword that could present a serious handicap in fluctuating environments. The authors of this counterintuitive idea have published only verbal models to support their claims.

Here we present the results of individual-based stochastic modelling of competition between two asexual species differing only by their evolvability. They show that, in changeable environments, less evolvable species could outperform their more evolvable competitors in a broad area of a parameter space, regardless of whether the conditions fluctuated periodically or aperiodically. Highly evolvable species prospered better nearly all the time; however, they sustained a higher probability of extinction during rare events of the rapid transient change of conditions.

Our results offer an explanation of why sexually reproducing species, with their reduced capacity to respond adaptively to local or temporal environmental changes, prevail in most eukaryotic taxa in nearly all biotopes on the surface of Earth. These species may suffer several important disadvantages in direct competitive battles with asexual species; however, they might win in changeable environments in the more important sorting-according-to-stability war.

## Background

Most of the eukaryotic organisms on Earth reproduce sexually, despite the existence of many obvious disadvantages, including two twofold costs of this complicated mode of reproduction (the cost of meiosis and the cost of males (Otto 2009). Many models and hypotheses have been published within the past 40 years to describe the conditions under which sexually reproducing organisms can outperform their asexual competitors, including asexual mutants (Bell 1982; Maynard Smith 1978). For many, but not all, of these models, see, for example the DNA repairing models (Horandl and Hadacek 2013) or variants of Muller ratchet models (Kondrashov 1982; Muller 1964), such conditions are relatively special and occur only in certain ecological situations. At the same time, sexual reproduction is predominant in all groups of eukaryotic organisms regardless of their taxonomic position or ecological strategy (Charlesworth 2006). Moreover, the obligate sex prevails in many taxa, despite the fact that theoretical studies clearly show that it is nearly always outperformed by the facultative sex, the condition-dependent alternation of many rounds of asexual reproduction with a round or rounds of sexual reproduction (Bell 1982; Green and Noakes 1995).

An interesting verbal model explaining the origin and persistence of sexual reproduction was suggested by Williams (1975), in his seminal book Sex and Evolution pp. 145-146, 149-154, 169. He argued that, paradoxically, sexual species can take advantage of their lower ability to evolve. Due to the effect of gene flow and due to the negative influence of segregation and recombination on the heritability of phenotypic traits and fitness, the ability of populations of sexual species to adapt to local environmental conditions is lower in comparison with populations of asexual species. Therefore, populations of sexual species usually retain a large part of their genetic polymorphism, including alleles that are suboptimal under present local conditions. Such alleles interfere with the ability of sexual species to fully and deeply adapt to local conditions. However, their presence can be extremely useful for the survival of the population and species when local conditions change.

It was also suggested recently that one of the important differences between asexual and sexual organisms is a much higher incidence of frequency-dependent selection, including selection in favor of heterozygotes, in the sexuals (Flegr 2010). Together with pleiotropy and epistasis, the presence of a certain amount (possibly not too high an amount) of alleles with such frequency dependent effects on fitness could stabilize the composition of the gene pool of a population, which would strongly decrease its ability to respond to directional selection. One of the implications of this theory is that sexual species are favored in randomly or periodically fluctuating environments, i.e., in most environments on the surface of Earth, due to their lower ability to evolutionarily respond to changes in their environment. The stabilization of the composition of the gene pool by frequency dependent selection not only prevents the population from the elimination of momentarily suboptimal alleles (Williams 1975) but also limits its ability to respond to selection, and by doing so, protects the population against an adaptation to transient changes in its environment (Flegr 2013).

The counterintuitive idea of Williams regarding the advantage of lower evolvability has been theoretically studied by several authors in the context of the origin and maintenance of sexual reproduction, for review see Toman and Flegr (2018) and (Kondrashov 1993). The models show that under special conditions (specific genetic architecture, alternating stabilizing and disruptive selection, etc.), sexually reproducing organisms can outperform their asexual competitors (Gandon and Otto 2007; Roughgarden 1991). For example, Roughgarden (1991) showed that, in fluctuating conditions, returns of frequencies of particular phenotypes to Hardy-Weinberg equilibrium in each generation bounds the variance in the mean fitness of sexual species, which automatically results in the higher geometrical mean of fitness, and therefore better performance of the sexual species in competition with the asexual species. However, neither competition of two asexual species or two sexual species differing only in evolvability, nor the character of environmental fluctuation favoring sexuality, has ever been studied in detail.

The aim of the present study is to test the validity of the verbal models of Williams (1975) and Flegr (2013) using a numeric individual-based stochastic model. Specifically, we searched for combinations of parameters under which a lower ability to adaptively respond to selection alone (not in combination with amphimixis) is advantageous and may result in the victory of a less evolvable species over its more evolvable competitor. Our study has been inspired by, and has implications for, theories of origin and sustaining of sexuality. However, it must be emphasized that we are going to study not the competition of sexual and asexual species but the competition between two asexual species: a more evolvable asexual species, which can freely respond to selection, and a less evolvable asexual species, whose members are penalized for deflecting from its original phenotype. This is because the aim of present study is not to test whether sexual species outperform their asexual competitors under fluctuating conditions. It is possible that under such conditions the sexual species could overrun its competitor not because of its lower evolvability but due to other already known advantages of sexuality, for example the capacity to select two or more positive mutations in parallel, the ability to stop Muller ratchet etc. In our model of competition of two asexual species, we intentionally stripped the less evolvable species of all other potential advantages to see whether, under certain conditions, just the lower evolvability could ensure its victory in a competition.

## Methods

### Model

We modeled the indirect competition in resistance toward extinction of two asexual species that differ only by their evolvability: the more evolvable plastic species and less evolvable elastic species. The terms “plastic” and “elastic,” used in the present paper to distinguish two species, have been borrowed from physics. In “plastic” organisms, the size of the change in population mean phenotype in response to constant force (constant selection pressure) is unceasing, regardless of how far the phenotype is from its original state. In the elastic species the size of the change in population mean phenotype in response to constant force is negatively proportional to the distance of the current phenotype from its original state. At a certain distance, this response can decrease to zero and the phenotype of the organisms stops responding to selection.

The time in our stochastic, individual-based model is discrete, i.e., all births and deaths and environmental changes happen simultaneously, and is measured in “seasons”. In each season individuals propagated with a given probability and died with another probability. Therefore, each season can be considered to be one reproduction period of a species. We used this model to study the competition of two species, plastic and elastic asexual species, living in an unstructured environment characterized by one (periodically or aperiodically fluctuating) environmental variable *E*_*env*_, e.g., temperature. In the periodically fluctuating environment, the current temperature, and at the same time the optimal temperature, for an organism to be adapted to, *E*_*env*_, is represented by a sinusoid that is characterized by its amplitude *A* and period *T* (for the sake of clarity, the amplitude is understand as (max-min)/2, not max-min; formally it is half the “peak to peak amplitude”). Fig. 1a shows the *E*_*env*_ curve (red line) for amplitude 1.3 and period 500. The aperiodic conditions are described by a stochastic curve with particular amplitude *A*, generated by randomly adding or subtracting the constant increment *ΔE* to/from *E*_*env*_ with probability *P*_*E*_ per season. Here, the rate of environmental change is characterized by the pseudoperiod *T* (determined by the combination of *ΔE* and *P*_*E*_), which is numerically equal to such a period *T* of the periodic model, for which the average speed of *E*_*opt*_ change from *-A* to *A* is the same for both models. The value of *ΔE* was fixed to 0.2 and *P*_*E*_ for each pseudoperiod was computed in advance by the Monte Carlo method. The value of *E*_*env*_ is bounded – when after an increment, it would exceed *A* (or fall below *- A*), it is reset to *A* (or *-A*). Fig. 2a shows a stochastic curve *E*_*env*_ (red line) with amplitude 1.2, *ΔE* = 0.2, and *P*_*E*_ = 0.2. For both periodical and aperiodical conditions, the changes of temperature are either continuous (the change of *E*_*env*_ immediately affects the organisms) or punctuational (the intrinsic continuous change of *E*_*env*_ manifests itself with probability *P*_*m*_ per season, see Figs. 1b and 2b for *P*_*m*_ = 0.1).

**Fig. 1.**
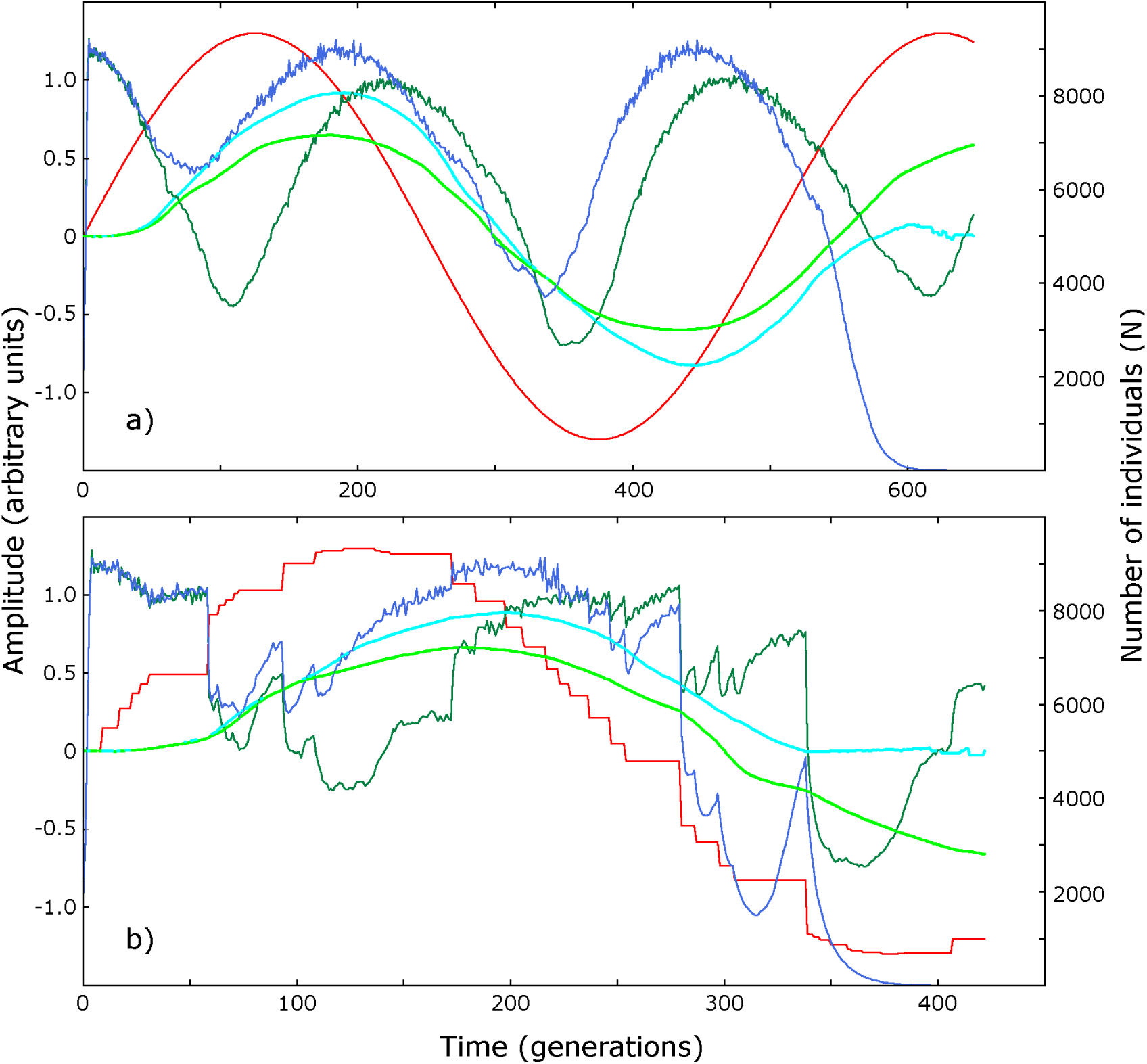
Competition of evolutionarily plastic and elastic species under periodically changing conditions. The conditions (the variable temperature - the red line) changes either continuously (the upper part a) or discontinuously (lower part b). The dark blue, dark green, turquoise and light lines show the size of plastic species, size of elastic species, mean phenotype (E) of elastic species and mean phenotype of plastic species, respectively.

**Fig. 2.**
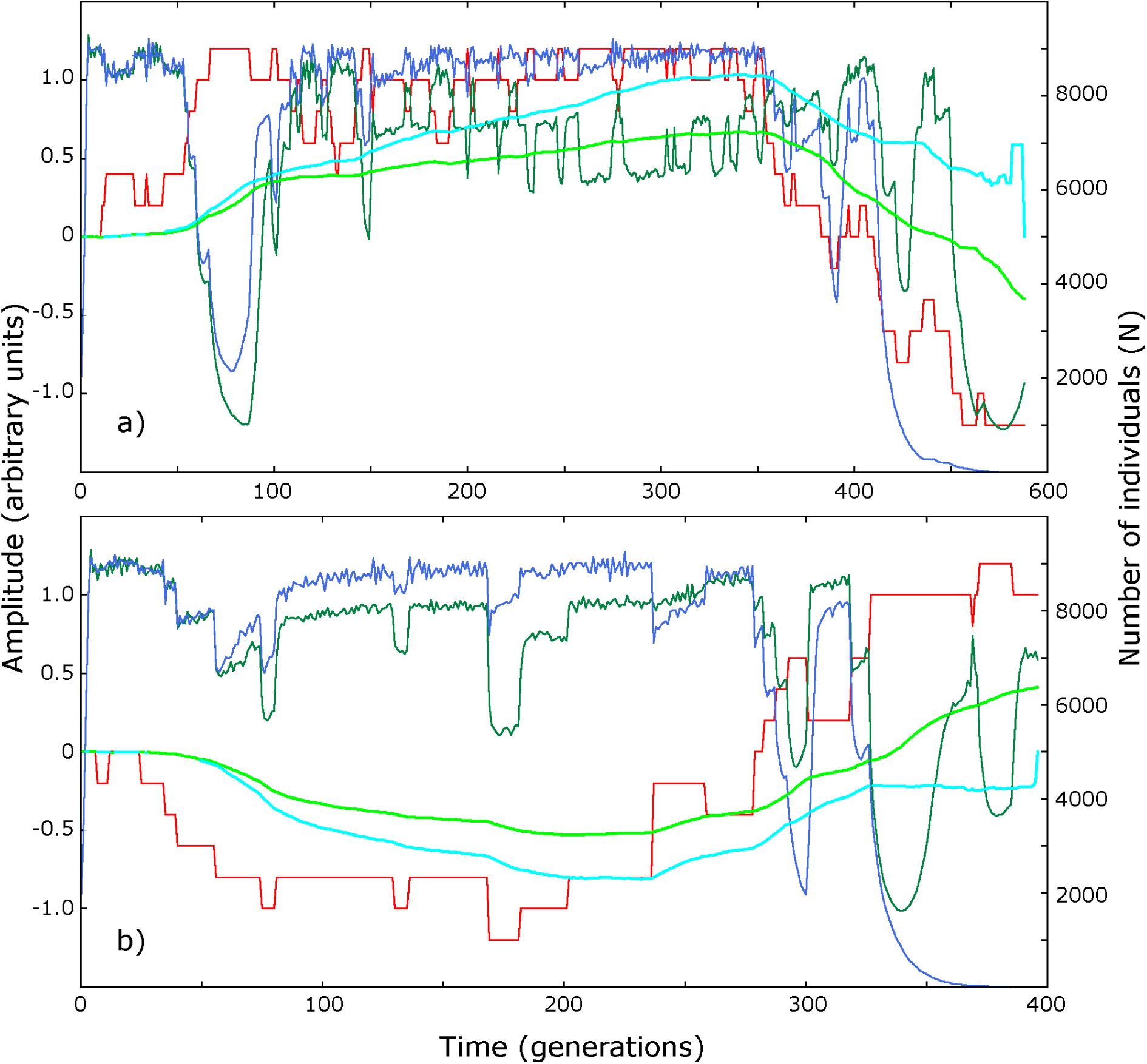
Competition of evolutionarily plastic and elastic species under aperiodically changing conditions. The conditions (the variable temperature - the red line) changes either continuously (the upper part a) or discontinuously (lower part b). The dark blue, dark green, turquoise and light lines show the size of plastic species, size of elastic species, mean phenotype (E) of elastic species and mean phenotype of plastic species, respectively.

The size of the populations of both plastic and elastic species are independently density-regulated by a turbidostatic mechanism (Flegr 1994). Namely, the probability of the death of an individual, *P*_*u*_, is *k*_4_ *N*^2^ + *k*_5_, where *k*_5_ is the probability of dying due to senescence or due to accident (density-independent component of mortality), *N* is the number of individuals of a particular species, and *k*_4_ is the probability of death due to a density-dependent process, e.g., due to contracting a directly transmitted parasite, the event probability of which increases with the square of *N* (Flegr 1997). In our simulation experiments, we set *k*_*4*_ = 5. 10^−9^ (which gives a maximum equilibrium population of approximately 10 000), and *k*_*5*_ = 0.1. The phenotype of each individual is characterized by a single parameter *E*, reflecting its body temperature (and therefore also indirectly the optimum temperature to live in). In each season, any individual can either die (with probability *P*_*u*_), reproduce (with probability *P*_*n*_), or do nothing. When a particular organism reproduces, its descendant either inherits the parental phenotype *E* or (with probability *P*_*x*_ = 0.1) mutates, i.e., its *E* increases or decreases by 0.05. The probability of reproduction of a member of plastic and elastic species is

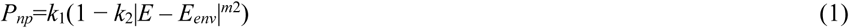

and

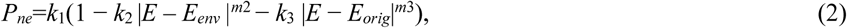

respectively. The probabilities of reproduction were bounded – when it fell below 0, it was reset to 0. *k*_*1*_ is the probability of reproduction under ideal conditions, i.e., when the temperature optimum of a particular individual corresponds to the actual temperature and this temperature corresponds to the temperature existing at the time of the origin of the species *E* = *E*_*env*_ = *E*_*orig*_. *k*_*2*_ and *m*_*2*_ characterize the penalty for the deviation of the phenotype of an individual from the current temperature *E*_env_, i.e., for *E* ≠ *E*_*env*_, and *k*_*3*_ and *m*_*3*_ characterize the penalty (paid by elastic species only) for the deviation of the current phenotype of an individual from the original phenotype (Flegr 2013). In our model, the *E*_*orig*_ = *E*_*env*_ = 0 at the start of our simulation experiment, and the penalty was positively correlated with the squared difference between *E* and *E*_*orig*_. The existence of this second penalty is the only difference between elastic and plastic species, and this part of the equation is responsible for the elastic nature of evolutionary responses of sexual species, i.e., for the slowing down and final stoppage of an evolutionary response of the elastic species on selection. In the presented simulations, we set *k*_*1*_=*k*_*2*_=1, *k*_*3*_=0.6, *m*_*2*_=*m*_*3*_=2 and for this combination of parameters the analytical solution of the equation (2) shows that the elastic species will stop responding to selection at the distance E = 0.625 E_env_ while the plastic species will continue responding until E = E_env_.

In the present model, the low evolvability of one of the competing species was ensured by introducing a penalty for the deviation of the phenotype of an individual from its original phenotype. We can imagine, for example, that adapting body temperature to a value that better corresponds to the new environmental conditions could decrease the amount of energy needed for thermoregulation. However, it could also impair the functions of thousands of enzymes adapted to the original body temperature.

### Implementation of the model

The model is programmed as a modular web application in the PHP language. The parameters are entered via a web form. At the start of the simulation, the time series of *E*_*env*_ is computed in AWK, the interpreted programming language. The computer time demanding part of the program, namely the individual-based simulation of population processes, are written in C. Numerical results are visualized using gnuplot. The web application for the simulation of competition that can show 1) the course of one simulation experiment and 2) aggregate results for N repeated simulation experiments performed with the same parameters is available at http://fyzika.ft.utb.cz/eng/index.php.

### Procedure

Three hundred individual simulation experiments were performed for each combination of amplitude and period (or pseudoperiod for aperiodic changes) (*A*: 20-980, step 60, *T*: 0.8-4.0). Each run was terminated after 10,000 seasons or when one of the species went extinct. The numbers of plastic species and elastic species extinctions was compared with a two-sided Pearson’s Chi-squared test (goodness of fit test). The parameter space was divided into four areas as shown in figures 3-7: the blue area where the evolutionarily elastic species wins significantly more often (p < 0.05), the red area where the evolutionarily plastic species wins significantly more often, the gray area where the difference in both species surviving was not significant, and the white area where both species usually survive until the end of the simulation experiment, i.e., for 10,000 seasons.

**Fig. 3.**
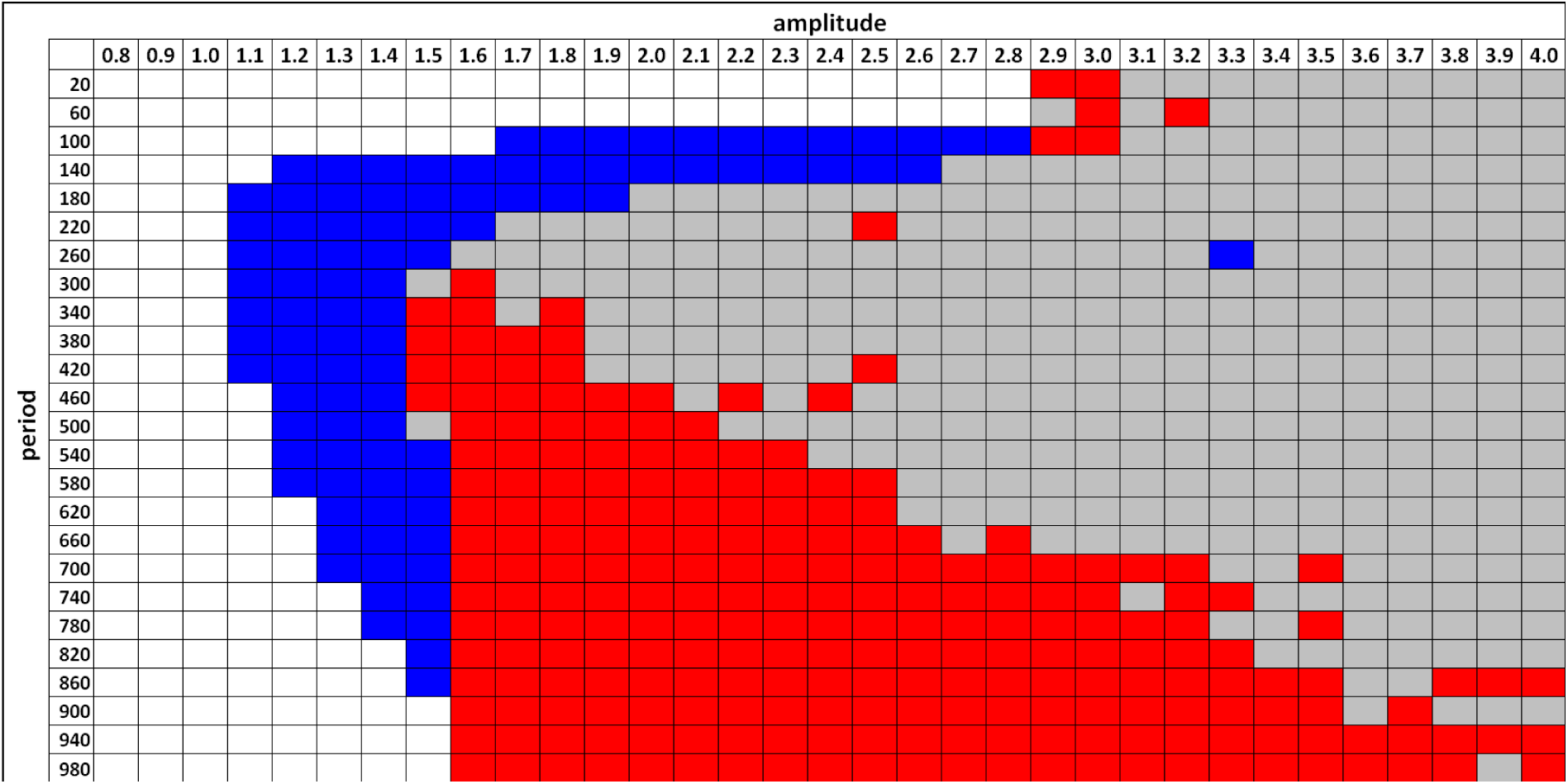
Competition of the plastic and elastic species in various parts of the parameter space under periodically and continuously changing conditions. The blue area denote combinations of amplitude and periods of environmental changes in which the evolutionarily elastic species win significantly more often (two sided goodness of fit test, p < 0.05), the red area the combinations in which the evolutionarily plastic species win significantly more often, the gray area the combinations in which the difference in survival for both species was not significant, and the white area the combinations in which both species usually survive until the end of simulation experiment, i.e. for 10,000 seasons.

## Results

We studied the extinction times of evolutionarily plastic and evolutionarily elastic organisms under conditions of both periodically (Fig. 1) and aperiodically (Fig. 2) changing environments. Under both conditions, the performance of the plastic species (measured by mean fitness or mean population size) was better most of time. However, the final result of the competition depended on the rate of the changes and the magnitude of the changes.

The results of the simulation for continuous periodic changes showed that under conditions of moderately-sized changes, or under conditions of rapid changes, the elastic species won significantly more often than the evolutionarily plastic species. On the other hand, the evolutionarily plastic species won when the changes were slow and the size of change was large, see Figs. 3-7. The evolutionarily plastic species also won in a second small region of the period-amplitude parameter space, namely for the periods 10-130 and the amplitudes 2.9-3.1, see Figure 7.

**Fig. 7.**
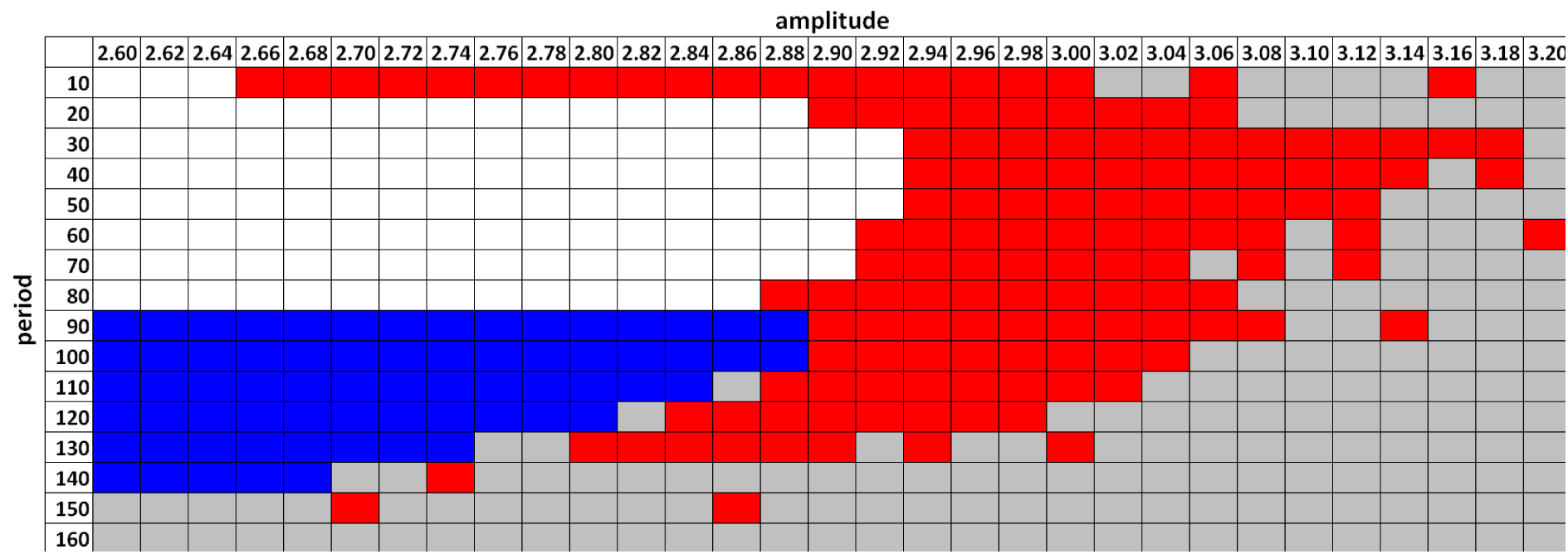
Competition of the plastic and elastic species in the short period-large amplitude region of the parameter space under periodically and continuously changing conditions. *For the legend see the Fig. 3*

For the combination of parameters used in our simulation, aperiodic conditions favored plastic species for amplitudes larger than 1.5, compare Fig. 3 and 5. In contrast, discontinuous changes (Fig. 4 and Fig. 6) somewhat favored the elastic species. The size of the elastic species-winning area of the period-amplitude parameter space was slightly larger and its position and shape differed (see the Discussion).

**Fig. 4.**
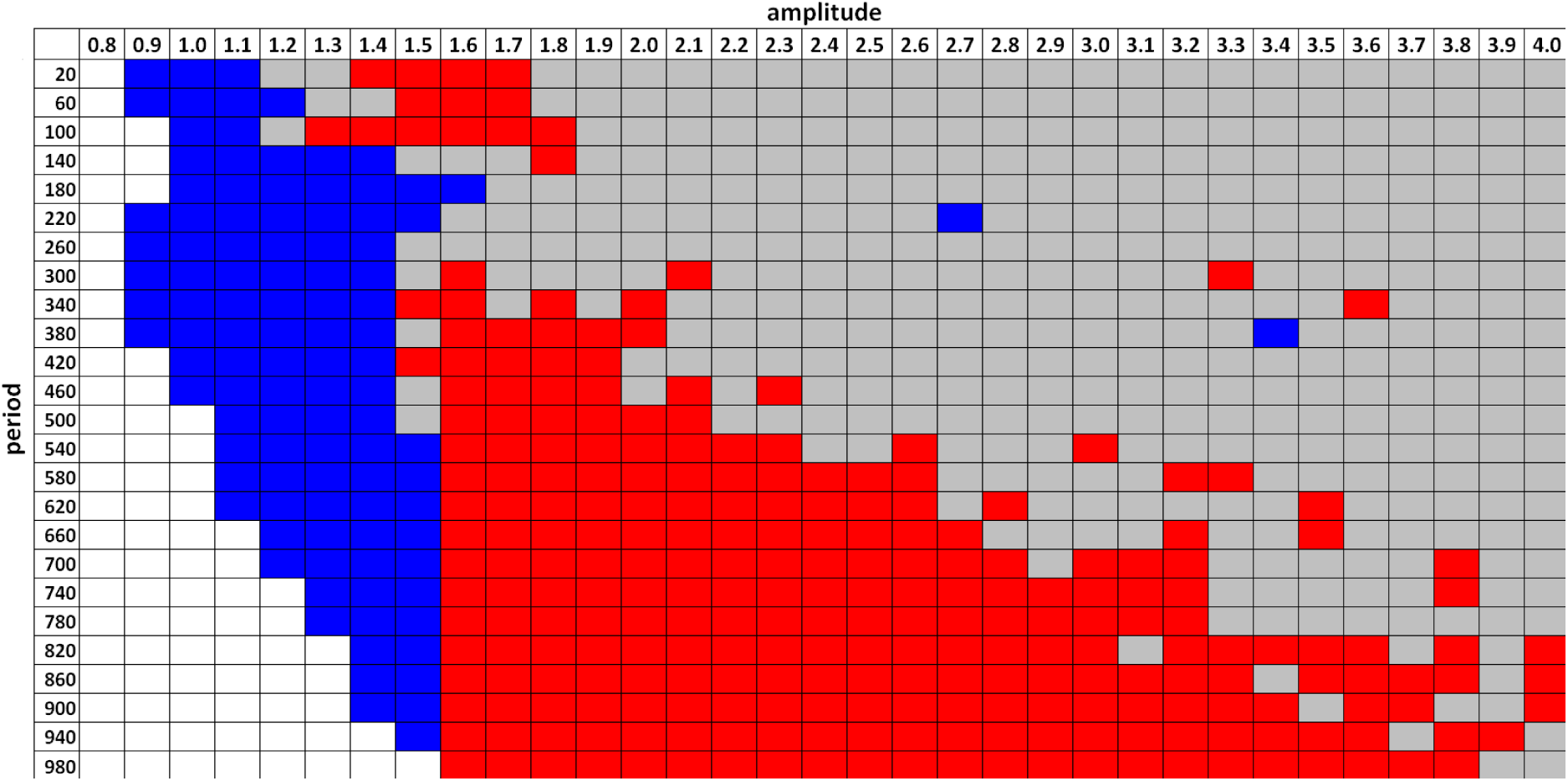
Competition of the plastic and elastic species in various parts of the parameter space under periodically and discontinuously changing conditions. *For the legend see the Fig. 3*

**Fig. 5.**
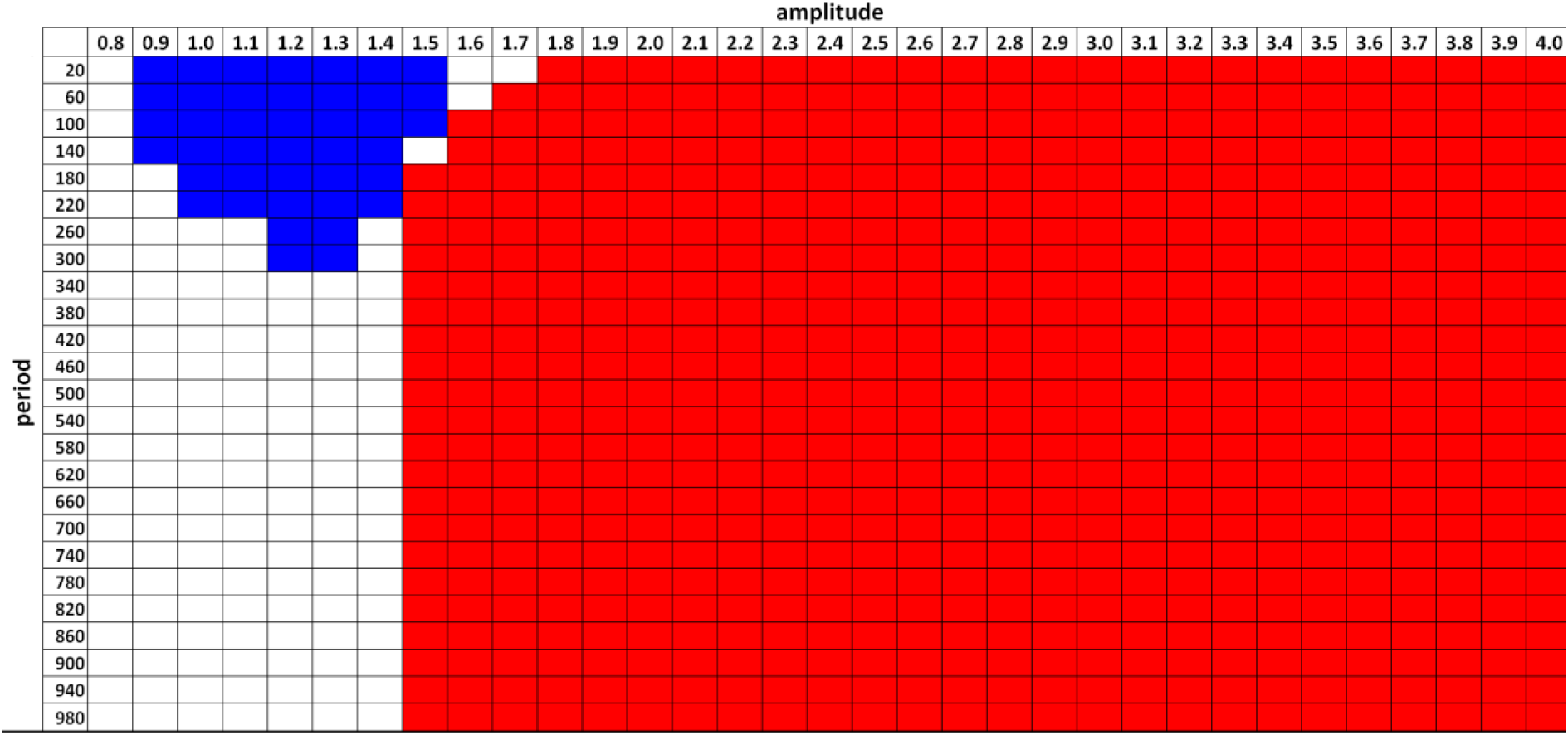
Competition of the plastic and elastic species in various parts of the parameter space under aperiodically and continuously changing conditions. *For the legend see the Fig. 3*

**Fig. 6.**
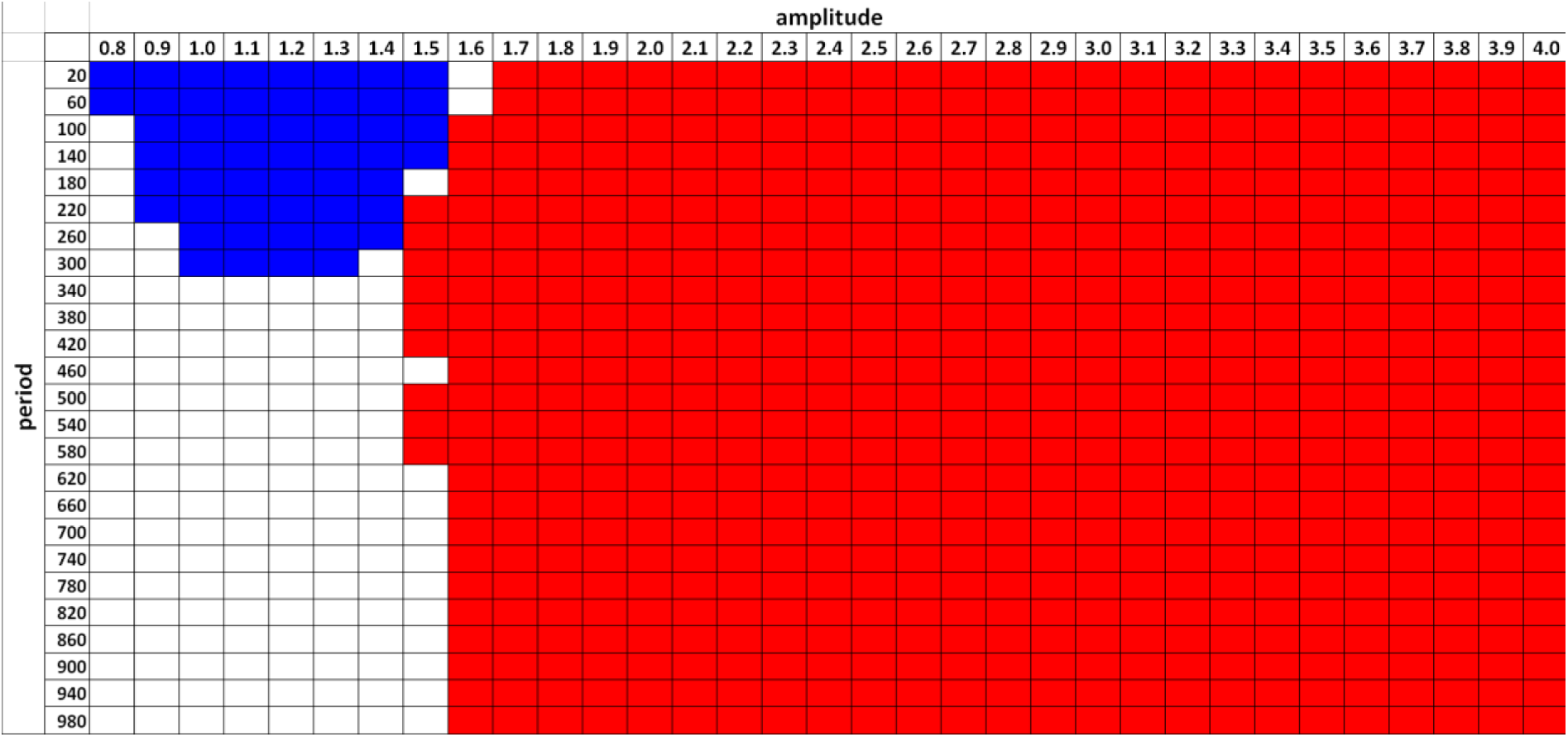
Competition of the plastic and elastic species in various parts of the parameter space under aperiodically and discontinuously changing conditions. *For the legend see the Fig. 3*

## Discussion

Our results confirmed that, under fluctuating environmental conditions, the evolutionary passivity of species with low evolvability, namely their limited ability to respond to selection by adaptive phenotypic change, could provide them with an advantage when indirectly competing (see below) with more evolvable species in a broad area of parameter space. Under such conditions, the evolutionarily passive species had a lower risk of extinction. This advantage was slightly higher when the environmental conditions fluctuated periodically and discontinuously.

At face value, this result might seem rather counterintuitive. In our model, the members of the evolutionarily passive (in our model elastic) species differed from the members of the evolutionarily plastic species only by the existence of a penalty that they had to pay for a deviation of their phenotype from the phenotype that they had at the start of each simulation run. Specifically, the size of the penalty (a decrease of the probability of reproduction in a particular time step) was directly proportional to the square of this deviation. Due to this term, the evolutionarily passive species responded to selection pressure elastically (at first easily, but slowing their response in time, stopping altogether in the end) and therefore only partially. It could adaptively respond to small changes in environmental conditions, but not to large ones.

The elastic species showed a lower risk of extinction than the plastic species in a part of parameter space (the blue area) because the population of the plastic species adapted from time to time to transiently changed conditions, not being able to readapt quickly enough when the conditions returned to or overshot the norm. On the contrary, the phenotype of members of evolutionarily elastic species did not deviate from their original phenotype too much. Carriers of “adaptive” mutations were rewarded for the phenotype that was better suited to their actual environment. However, at the same time, they were penalized for the deviation of their phenotype from the phenotype that they had had at the beginning of the simulation runs. The sharp boundary between the blue and red areas around the amplitude 1.5 existed for all but very quickly periodically changing environments because the elastic species can survive only for a very short time once *E* minus *E*_*orig*_ is greater than about 1.5 – for larger differences, the mean birth rate is always smaller than the mean death rate. When the environmental changes were periodical and continuous, the elastic species outperformed the plastic species in a broad interval of rate of environmental change (periods 100-860) when the size of environmental changes was relatively small (amplitude 1.1-1.5), and also when the size of environmental changes were moderate and large (the amplitude 1.5-2.8) and the rate of changes was large, namely the period was in a relatively narrow interval 100-180. In the later part of the elastic species-winning area (the blue high periodicity tail) the mean phenotype of the plastic species increased or decreased strongly and settled down rather close to one of the boundaries of the fluctuation interval. In contrast, the mean phenotype of the elastic species did not change and remained close to *E*_*orig*_. In consequence, the plastic species had slightly larger fluctuations in abundance, which sooner or later led to its exinction. Under conditions of aperiodic and also discontinuous changes, the blue high periodicity tail of the elastic species-winning area was absent. In the white area of no statistical difference above the blue tail, the fluctuations of the plastic species were too small to cause exinction, so both species survived. In the grey area, both species went extinct very quickly and did not survive the first environmental fluctuation. The evolutionarily plastic species usually won in slowly changing environments, especially when the changes were large (amplitudes > 1.6). In periodically fluctuating environments, the amplitude that was most favorable for the plastic species was about 1.6, and the resistance to the increase of the amplitude raised with the size of the period, i.e., the plastic species significantly outperformed the elastic species when the rate of change was slow enough, e.g., when the period was at least 980, even when the amplitude was as large as 4. Optically, the size of the main part of the plastic species-winning area is large. However, it must be emphasized that in this region both species usually go extinct during the first period of environmental change (the elastic species earlier). It is therefore questionable whether this combination of parameters is biologically relevant. The plastic species also won when the changes were very fast (the period or pseudoperiod was 10-110) and the size of changes was large but not the maximum (amplitude 2.9-3.1), Fig. 7. Under these conditions, the carriers of adaptive mutations outperformed other members of plastic (and also elastic) species; however, the number of carriers of standard phenotypes remained relatively high at the moments when the environmental conditions returned. This probably saved the plastic species from extinction. In the red bulk of this smaller part of the plastic species-wining area, the phenotypes of both species were close to *E*_*orig*_. Both populations declined rapidly and fluctuated at a low level. Typically, the population of the plastic species was a bit smaller and vanished first. In the left-sided tail of the red bulk (e.g. for *A* = 2.7, *T* = 10), mean phenotypes of both species varied very little. Population sizes both increased and then decreased in synchrony, but the elastic species usually went extinct a little bit earlier than the plastic one. This likely occurred when the mean phenotype of the elastic species finally changed a bit, either due to selection or due to drift when the population size decreased to a very low value. When the conditions fluctuated aperiodically, the amplitude most favorable for the plastic species was > 1.8; for these amplitudes, the plastic species outperformed the elastic species even when the rate of change was very large (pseudoperiod > 20).

During all simulations, the evolutionarily plastic species outperformed and therefore outnumbered the elastic species most of the time. However, in rarely occurring situations, e.g., when the conditions changed unusually strongly and rapidly in a non-periodically fluctuating environment or, in a periodically fluctuating environment, when many adaptive mutants appeared unusually early during the selection period and therefore shifted the mean phenotype too far from the norm, the plastic species was reduced to zero or to a very small value. In very small populations, genetic drift (i.e., chance), rather than fitness, determines the destiny of individuals. Also, the number of arising mutations is too low there. Therefore, any small population, including the population of plastic species, loses the ability to adaptively respond to changes in its environment.

It is important to emphasize that we modeled the indirect competition of two species that did not interact ecologically, for example, two species that did not exploit any common resource or that lived in separate areas of the environment. When, for any reason, the population of the first species increased (decreased), the situation of the second species was not influenced by this. Therefore, the subjects of this study are macroevolutionary or macroecological phenomena, namely the sorting of species or populations on the basis of stability (stability-based sorting), rather than microecological phenomena like direct interspecific or intraspecific competition for resources. In principle, we modeled a situation analogous to a scenario in which plastic and elastic species are introduced 300 times to two identical isolated islands and then counted how many times each species survived longer on its own island. If direct competition was permitted, e.g., when growth of the populations of both species is affected by the same parasite (*P*_*u*_ = *k*_*4*_ *(N*_*p*_ *+ N*_*e*_)^*2*^), the result of our simulation was different. Under such conditions, the plastic species outperformed the elastic species in the whole parameter space (results not shown). However, competition without any direct ecological interaction operates in many groups of organisms. For example, genetically different lineages of parasites as well as different parasitic species rarely meet in one host even in situations where when they live in the same area (Morand et al. 1999). The same holds for species that exploit various temporary habitats like forest openings, puddles, rotting fruits or animal and plant remains. Indirect competition, however, also plays an extremely important role in species with “normal” ecology. Over long timescales, most habitats on Earth are unstable. Particular localities come and go, old localities turn uninhabitable for particular species and new inhabitable localities originate. When a species colonizes a new suitable locality, its population is at least transiently liberated from its competitors. Frequently, on long-term time scales, the species that are weak direct competitors can win when they are able to quickly colonize new suitable locations and there produce many new colonists before their stronger competitors arrive and outcompete them, or before their locations cease to exist. Actually, it can be speculated that the low growth rate of weak competitors could be the very reason for their final victory because it can help them to escape overexploitation of their resources, which can help them to keep their environment (e.g. the host organism in the case of parasitic species) inhabitable for a longer time.

When not only rapid fluctuations but also some slow and systematic (unidirectional) change occurs in a certain environment, and when the plastic species succeeds in surviving the fluctuations long enough, the plastic species would finally win over its elastic competitor. The penalty paid by the elastic species for its out-of-date phenotype grows with the systematic change of the environment until it turns incompatible with the survival of the species. Before it happens, however, the elastic species could speciate, and the new species could transiently turn plastic and therefore acquire the ability to adapt to changed conditions as was proposed in some punctuational theories of evolution (Carson 1968; Flegr 2010; Mayr 1954; Templeton 2008). After such an “evolutionary reset”, the new species returns to elasticity (by the slow accumulation of genetic polymorphism, especially by the accumulation of mutations with a frequency-dependent effect on fitness). The new elastic species would probably outcompete the old and obsolete elastic species (Pearson 1998), and the competition between the plastic species and new elastic species can continue (Flegr 2013).

Seemingly, our results contrast with results of comparative studies that mostly show that asexual species are more common in unstable environments (e.g. disturbed habitats, intertidal, temporal pools, temperate habitats) while sexual species are relatively more common in stable environments, e.g. in tropics (Bell 1982). It is necessary to reiterate once more that we studied the competition of a more evolvable asexual species with a less evaluable one, not the competition of asexual with sexual species. Also, we did not compare the performance of these two species in more stable and less stable environments, but in environments that differed by the nature of their changes – by the combination of their amplitudes and frequencies. The figures 3-7 clearly show that it is not possible to say which species is better adapted to rapidly or slowly fluctuating environments – it always depends on the amplitude of the environmental change. Finally, yet importantly, the results of recent comparative studies performed exclusively on ancient asexuals, the species that successfully passed the test of long-term (more than 1 million years) persistence in nature, showed that whenever the environment of asexual species differed from the environment of its related sexual species, the asexual species lived in a less heterogeneous environment.

In comparison with real systems, our model favors the plastic species in two important ways. First, in real organisms, the fitness of an individual is determined by several traits rather than just one, as it is in our model. Moreover, each trait is usually determined or influenced by many genes, the effects of which are often not additive (Griffiths and Neumann-Held 1999). In such a multidimensional adaptive landscape, the rapid adaptation of plastic species to the drastic (rapid and large) changes of an environment is probably much more difficult than in the unidimensional adaptive landscape that is the subject of our simulations. The difficulty of quick return to the original phenotype probably grows with the number of dimensions, and it is even possible that the plastic species could finish trapped, or at least transiently trapped, in a certain location of the adaptive landscape (Schwartz 2002).

Second, in real systems, the evolutionary passivity and elasticity of species is expected to be the consequence of their sexual reproduction. Therefore, in sexual species, evolutionary passivity is accompanied by the persistence of a large amount of genetic polymorphism that can be sustained in the population by various mechanisms related to sex (Burger 1999; Roughgarden 1991; Waxman and Peck 1999). For example, in sexual species, the fixation of genotypes adapted to local conditions and the extinction of locally maladapted genotypes is very slow or even impossible due to segregation and recombination, as well as due to gene flow, the hybridization of members of a local population with migrants (Dias and Blondel 1996; Haldane 1956). Moreover, on a large time-scale, sexual species can sustain their diploid status, while diploid or polyploid asexuals will finely turn functionally haploid due to the accumulation of mutations in spared copies of genes (Lewis and Wolpert 1979). Due to their diploidy, they can maintain high genetic polymorphism in their gene pool by the heterozygote advantage effect, which represents a special type of frequency dependent selection. In our model, both plastic and elastic species reproduce asexually, and thus they have comparable amounts of genetic polymorphism in their gene pools (as shown by the similar standard deviation of *E* in outputs of our program). Therefore, the elastic species in our model is deprived of its largest advantage – the ability to very quickly (although only transiently and only imperfectly) respond to rapid changes by the shifting frequencies of already existing (old) alleles. In contrast to a real situation, the rate of response to changes is similar in elastic and plastic species at the beginning of our simulations (as it was mostly fueled by rare mutations) and slows down in the elastic species as its phenotype declines from its original state. Nevertheless, our present results suggest that even when elastic species are stripped of this crucial advantage, they can outcompete the plastic species in a broad area of parameter space. Evolutionary passivity alone, without the usually accompanying higher polymorphism resulting in quicker evolutionary response to new selective pressure, can therefore explain the superiority of sexual species under fluctuating environmental conditions.

Typically, a positive correlation between the probability of local extinction and global extinction exists (Payne and Finnegan 2007). Therefore, a decreased probability of extinction in a fluctuating environment could be advantageous not only on an ecological time scale, but also in macroevolution. In agreement with the verbal arguments of G.C. Williams (1975), our results show that a possible reason for the long-term success of sexual species may be, paradoxically, their lower evolutionary plasticity (lower evolvability), which reduces the risk of extinction of the population or species in an environment with randomly or periodically fluctuating conditions. This means that sexual reproduction might not be the evolutionary adaptation that increases some aspect of direct or inclusive fitness of its carriers, as is suggested by most present theories on the origin of sex. Sex could rather be the evolutionary exaptation (Gould 2002; Gould and Lewontin 1979) that increases the chances of a given species and evolutionary lineages in the process of stability-based sorting and in the process of species selection (Toman and Flegr 2017; Vrba and Gould 1986).

## Authors’ Contributions

JF designed the study and wrote the manuscript, PP wrote the program, run the simulations and participated on writing the article. Both authors gave final approval for publication.

## Acknowledgements

Computational resources were provided by the CESNET LM2015042 and the CERIT Scientific Cloud LM2015085, provided under the programme “Projects of Large Research, Development, and Innovations Infrastructures. We would like to thank Charlie Lotterman for his help with the final version of the paper, and three anonymous referees and the editor of Axios Review Society for their extremely useful suggestions.

